# Focused acoustic-radiation-force microscopy for live-cell selective nucleus deformation and nucleus viscoelasticity measurement

**DOI:** 10.1101/2025.10.23.681068

**Authors:** Natsumi Fujiwara, Hiroki Okita, Midori Uno, Kanta Adachi, Kichitaro Nakajima, Nobutomo Nakamura, Mee-Hae Kim, Masahiro Kino-oka, Kotaro Ogawa, Kensuke Ikenaka, Hirotsugu Ogi

## Abstract

Acoustic-radiation-force (ARF) microscopy was developed for non-invasive, selective mechanical stimulation to a single live-cell nucleus and long-term measurement of its mechanical properties. High-frequency focused ultrasound wave achieved a spatially confined ARF field around target nucleus, and its cyclic deformation was induced by amplitude modulation on the ultrasound wave, which was captured by a dark-field microscopy. This method was applied to human mesenchymal stem cells, revealing their extremely high mechanical homeostasis.

---

Cell nucleus can perceive mechanical cues and respond by modulating its own mechanical properties to maintain cellular integrity, leading to downstream alterations in cellular functions. Numerous studies have aimed to elucidate cellular responses to mechanical stimuli by investigating the mechanical properties of the nucleus^1-3^. However, as summarized in **Supplementary Table 1**, no existing method can non-invasively and selectively deliver mechanical stimulation exclusively to a single live-cell nucleus while simultaneously enabling long-term measurement of mechanical properties of the nucleus.

Here, we develop an amplitude-modulated focused acoustic-radiation-force (ARF) microscopy (Fig. 1(a)) that enables selective, non-invasive deformation of nucleus, thereby overcoming the limitations of existing approaches. By generating a spatially confined ARF field around the nucleus region using an ∼80 MHz carrier wave amplitude-modulated at 0.5 Hz, its cyclic deformation of 1 Hz was induced, which was detected by a dark-field microscopy (Fig. 1(b), **Supplementary Movie S1**). It also allows long-term measurement for the same live-cell under physiologically relevant conditions. By monitoring phase lag and the strain of the nucleus deformation relative to ARF, we evaluated nucleus viscoelasticity in-situ (Figs. 1(c) and (d)).

**Figure 1.**
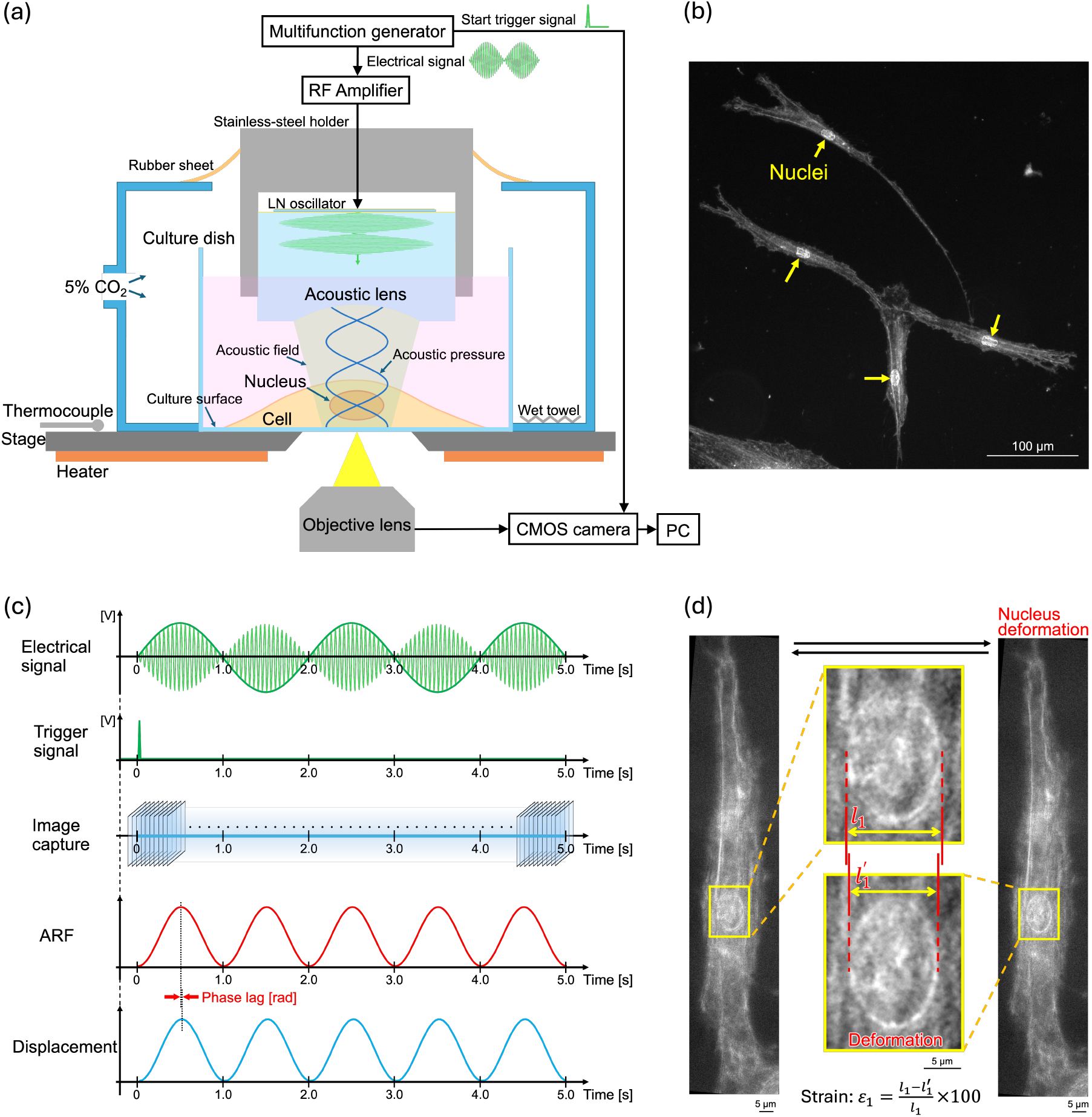
(a) Schematic of the experimental setup of ARF microscopy. (b) An image of live cultured cells acquired without labeling using a dark-field microscope located beneath the culture dish. (c) Schematic diagram showing measurement timing relationships. Top: Electric waveform sent to the LN oscillator. Second: Its trigger signal. Third: Image acquisition by the CMOS activated by the trigger signal. Fourth: Cyclic ARF synchronized with the trigger signal. Last: Deformation of the cell nucleus, showing a phase lag from the ARF change. (d) Determination of the in-plane strain.

The measurements were performed on randomly selected spindle-shaped cell based on the timeline shown in Fig. 2(a). The results are shown in Figs. 2(b)-(e). The phase lag of the nucleus is inherently smaller (cell 1 in Fig. 2(c)), but it becomes larger when connected to well-developed cytoskeletons (cells 3 and 4, Figs. 2(b) and (d)). This behavior is consistent with the highly viscous nature of the cytoskeleton, arising from dynamic processes such as polymerization and depolymerization with characteristic relaxation times of ∼1 s^4, 5^. Concerning cell 2, the phase lag increased after one minute, which may be due to its connection to the surrounding cytoskeleton.

**Figure 2.**
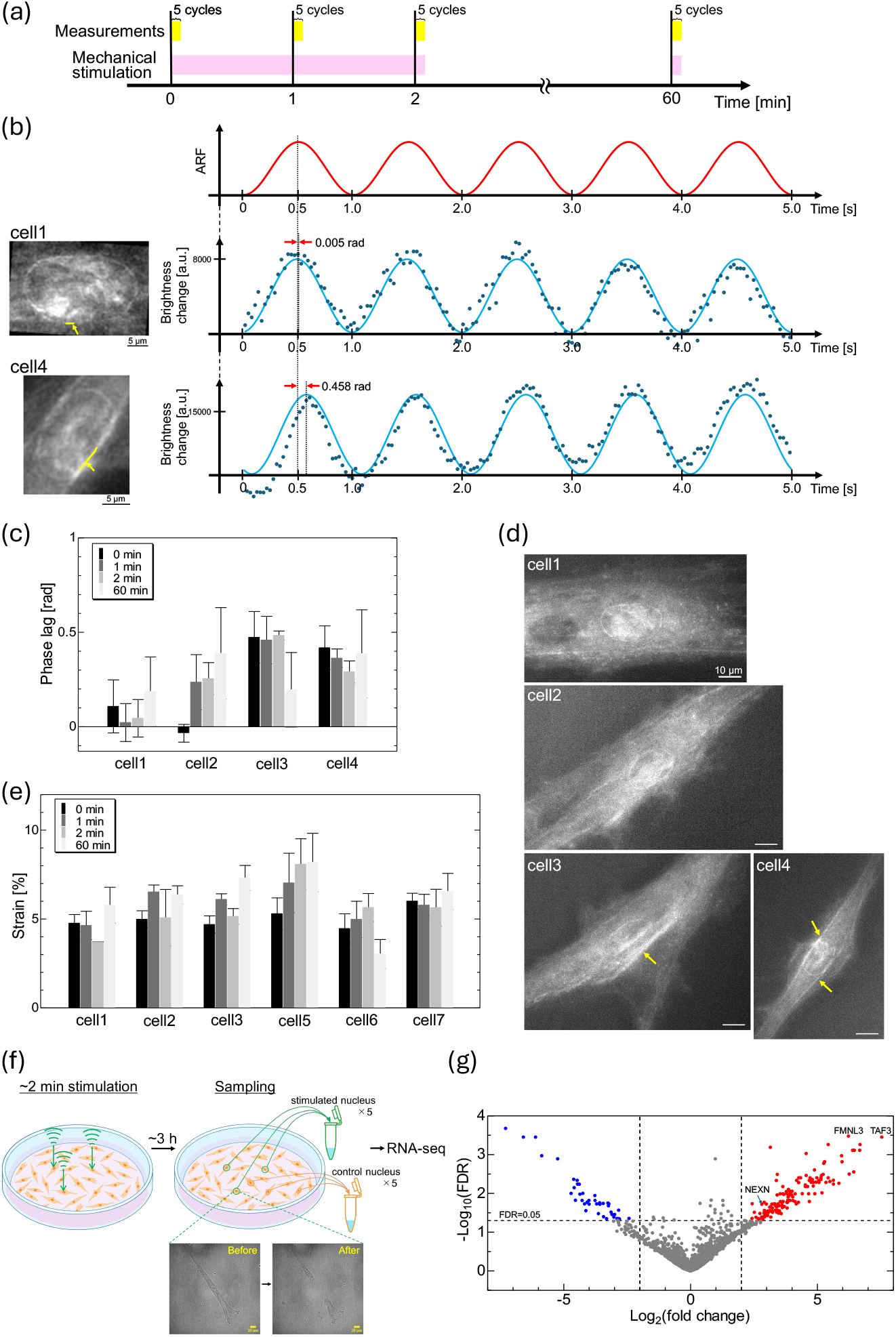
(a) Experimental timeline for mechanical stimulation and measurement of nuclear mechanical properties. (b) Changes in average brightness of pixels (dots) along nuclear envelope for nuclei lacking (middle) or possessing (lower) well-developed cytoskeletal connections. Red line shows the cyclic ARF and light-blue lines are fitted sinusoidal functions, from which the phase lag were extracted. (c) Phase lag values measured for four representative cells and (d) corresponding dark-field microscopy images. Yellow arrows indicate cytoskeletal structures connected to the nucleus. (e) Time-dependent changes in in-plane strain of nucleus. Cells 1-3 are the same as those in (d). (f) Schematic for gene expression analysis and representative images before and after the nucleus aspiration. (g) Volcano plot for the differential-expression analysis using DESeq2. Some genes, which are expected to contribute to mechanical homeostasis of living cells, are indicated.

While numerous studies have reported dramatic changes in nuclear mechanics by mechanical stimuli^6-9^, no significant time-dependent variation was observed in the strain in this study (Fig. 2(e). To our knowledge, this is the first demonstration of mechanical stimulation applied exclusively to the nucleus of a single live-cell in combination with long-term mechanical measurements. This finding suggests that the nucleus actively maintains mechanical homeostasis to protect chromatin from external perturbations; mechanical properties of a live-cell nucleus remain nearly unaltered.

A crucial advantage of ARF microscopy is the ability to establish both stimulated and unstimulated (control) cells within the same culture environment. Even under identical culture conditions and using the same type of culture dish, cell characteristics can vary between dishes^10^. Thus, stimulated and control cells should be compared within the same culture dish to accurately assess the effect of the stimulus. Our technique achieves nucleus-specific stimulation in single living cells, yielding robust and reproducible data (Fig. 2(f)).

Although no major change in the nuclear stiffness was observed upon stimulation (Fig. 2(e)), we performed RNA sequencing to assess potential molecular responses. Figure 2(g) shows the volcano plot, revealing significant upregulation of various genes, including those related to mechanical homeostasis such as formin-like 3 (FMNL3)^11^ and nexilin (NEXN)^12^, which promote polymerization and stabilization of actin filaments. Also, very significant upregulation is found for TATA-box binding protein associated factor 3 (TAF3), which participates in the formation of the transcription initiation complex by interacting H3K4me3 of chromatin^13^ and will alter the mechanical properties of chromatin, affecting the nucleus rigidity. The mechanical homeostasis is thus expected to be dynamically achieved in living cells.

In summary, we present the first method that enables non-contact, label-free, and nucleus-specific mechanical stimulation together with direct measurement of nucleus viscoelasticity in living cells. This approach provides a powerful framework for uncovering how cells sense and adapt to mechanical stress at the nuclear level. Notably, whereas previous studies reported substantial changes in nuclear mechanics following stimulation, our results revealed no such alterations, highlighting the remarkable mechanical homeostasis of living cells.

## Supporting information

Supplementary Note, Supplementary Table, Supplementary References, Descriptions of Supplementary Movies

## Methods

### Cell preparation

Cryopreserved human mesenchymal stem cells (MSCs) from a bone marrow origin (Lot no. 23TL253285; Lonza, Switzerland) were thawed and cultured according to the supplier’s instruction. They were cultured on a tissue culture-treated polystyrene surface (430166; Corning Inc., USA) in an MSC growth medium (PT-3001; Lonza) containing 1% antibiotic-antimycotic (15240062; Thermo Fisher Scientific, USA) at 37 °C in humidified air with 5% CO2. When cultures reached 70% confluence, the cells were detached with enzymatic treatment with 0.1% trypsin/0.02% ethylenediaminetetraacetic acid solution and reseeded at 1.5 × 10^3^ cells/cm^2^ on the plastic bottom with the tissue culture-treated surface (ib81156; ibidi GmbH, Germany). 24 h after the cell seeding, they were used for experiments.

### Development of acoustic-radiation-force microscopy

The laboratory made transducer can provide high-intensity ultrasound capable of generating ARF. The transducer consists of a quartz glass rod with an acoustic lens on its bottom surface (Extended Data Fig. 1(a)). An Au/Cr electrode was deposited on the top surface, onto which a 40-µm thick lithium niobate (LN) oscillator was bonded with epoxy resin. The Au/Cr electrode was then deposited on the upper surface of the LN oscillator. A cross-sectional shape of the acoustic lens portion is shown in Extended Data Fig. 1(b). The assembled ultrasound transducer was mounted in a stainless-steel holder, enabling the electric excitation via contact probes contacting the LN surface (Extended Data Fig.1 (c)). Because no damper material was attached to the LN oscillator, it can be oscillated in its fundamental resonance mode, providing high-intensity and long tone-burst ultrasound capable of generating ARF.

Because of the fixed-end reflection, an acoustic pressure node is formed at a height of one-quarter wavelength above the dish. The resulting ARF drives the nucleus elements toward this node, inducing deformation both along the vertical axis and within the plane through the Poisson effect (Extended Data Fig. 1(d)), the latter of which can be detected by the dark-field microscopy. The amplitude modulation onto the carrier wave was used to generate periodic ARF, causing the periodic deformation only for the target nucleus (**Supplementary Movie S2)**.

The electrical signal generated by the multifunction generator (WF1968; NF Corporation, Japan) was amplified by a high-frequency signal amplifier (P43A-20M1G; Insight K.K., Japan) and delivered to the transducer. The frequency of the longitudinal wave was between 80 and 90 MHz. To evaluate the focusing ability of the acoustic lens, we constructed spectroscopic acoustic absorption images as previously reported^14,15^ and obtained clear images of the cells (Extended Data Fig. 1(e)). In addition, irradiation of a plastic surface with extremely high-power ultrasound produced localized alterations at the focal point due to ultrasonic friction, allowing the focal position to be visualized (inset in Extended Data Fig. 1(e)). These observations indicate that the focal spot diameter is approximately 18 µm, matching the nucleus size.

The chamber was maintained at a culture environment of 5% CO2, 37 °C temperature, and 100% humidity. An objective lens was positioned beneath a 35-mm culture dish for dark-field microscopy, which monitors cyclic deformation of the nucleus without any labeling.

The only potentially invasive aspect of this method would be the temperature rise caused by the focused ultrasound. However, through the considerations outlined in **Supplementary Note**, we find that this is insignificant.

### Visualization of cross-sectional movement of nucleus by confocal microscopy

We observed nucleus deformation as projective images. However, the acoustic beam was expected to exert an ARF along the *z*-axis that would induce translational motion in the same direction as well. To investigate this *z*-direction motion, we observed the deformation behavior of the cross-section using confocal microscopy.

We obtained the cross-section images using a 100× objective lens (MRD71970; Nikon Engineering Co., Ltd., Japan). The nucleus was stained using NucSpot^TM^ Live 488 (40081; Biotium, Inc., USA). The dye was added to the culture medium at a concentration of one part per thousand, and the cells were incubated at 37 °C for ∼3 h. Just before capturing images, the culture medium was replaced with phosphate-buffered saline, and cross-sectional images were acquired. The in-plane cross-section images were captured at each optical section for four modulation cycles at 15 fps, synchronized to the modulation trigger. In total, 65 sections were recorded at 0.2 µm *z* -steps (260 deformation cycles) and subsequently corresponding x-z images were reconstructed.

The results reveal that both out-of-plane and in-plane expansion and contraction deformations are visible. In some cases, coexistence of translational and extensional motions are observed, but after removing translational motion, we found that the nucleus itself underwent cyclic expansion-contraction deformation. Therefore, they demonstrate that our system enables non-invasive induction of periodic nucleus deformation.

### Determination of acoustic pressure using silicon cantilever deformation

When a particle with radius *a*, mass density *ρ*_p_, and compressibility *κ*_p_ is located in a region where an acoustic plane wave propagates, the ARF acting on the particle parallel to the propagation direction, *F*_*z*_, is given by Eq. (1)^16, 17^,

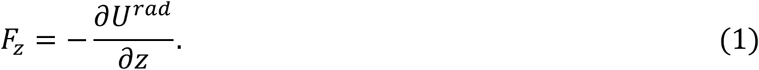

Here,

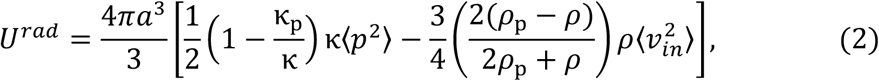

where *κ* and ρ denote the compressibility and mass density of the culture medium, respectively. ⟨*p*^2^⟩ and 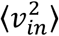 indicate the mean-square acoustic pressure in the culture medium and the mean-square velocity of the acoustic field at the particle position, respectively. Assuming one-dimensional ultrasound pressure *p* = *P*_*ac*_sin (*ωt* − *kz*) with pressure amplitude of *P*_*ac*_, the acoustic wave at the dish surface is expected to undergo nearly fixed-end reflection, and the sound pressure field of the composite wave is expressed as *p*_*tot*_ = 2*P*_*ac*_sin(*ωt*)cos (*kz*) (*z* is set to be zero at the dish surface). The maximum magnitude of the acoustic radiation force |*F*_*z*_| is then given by Eq. (3) using the relationship between the velocity and pressure as *v* = −(2*P*_*ac*_/*ρc*)cos(*ωt*)sin (*kz*):

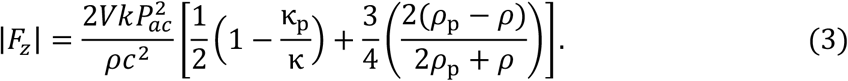

Where *V* is the nucleus volume, and *k* and *c* represent the wave number and sound velocity of the acoustic wave in the culture medium, respectively. The magnitude of the acoustic radiation force can be thus estimated when the pressure amplitude is known.

We evaluated the acoustic pressure amplitude *P*_*ac*_ and then the ARF using a silicon cantilever (ARROW-CONT-10; NanoWorld, Switzerland) set in the culture medium, whose stiffness is known: We focused the acoustic wave near the edge of the cantilever with a 45 *μ*m width and 450 *μ*m length. In this case, the ultrasound reflected at the cantilever surface induces the cantilever bending deformation through the interfacial acoustic radiation pressure *P*_*int*_ expressed below

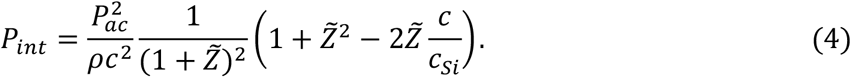

Here, *c*_*si*_ and 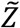 denote the sound velocity of the cantilever and the ratio of the acoustic impedance of culture medium to that of the cantilever, respectively. (Using the acoustic parameters for culture medium and Si, we have 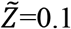.) The cantilever underwent the bending vibration by the modulated ARF. Because the brightness distribution along the cantilever reflects its deformation profile, we quantified the bending-vibration amplitude induced by the interfacial ARF by comparing the brightness distribution with the static cantilever-deformation experiments (calibration), where the cantilever was displaced in 1 µm increments using a microneedle (Extended Data Fig. 2(a)). The result is shown in Extended Data Fig. 2(b). The deformation amplitude was determined from the brightness at the inflection point of the brightness-displacement curve. For example, the experiment shown in Extended Data Fig. 2(b) yielded a deformation of 5.73 µm when 15 Vpp was applied to the transducer, corresponding to a force of 1.15 µN given by the spring constant of the cantilever (0.2 N/m). From the focal spot diameter of 18 µm (inset in Extended Data Fig. 1(e)), we obtain *P*_*int*_ = 4.50 kPa. *P*_*ac*_ is then estimated to be 3.96 MPa. Therefore, the maximum ARF force is estimated to be 400 nN in this case, using *c*=1530 m/s^18^, *a*=5 µm, 1 − *κ*_p_/*κ* = 0.2^17^ with an 85.5 MHz carrier wave. The input voltage can be changeable between 0 and 189.5 Vpp, indicating that our system has a very large dynamic range for ARF from 0 to 5.07 µN.

### Determination of phase lag and strain of nucleus deformation

The key feature of our system is the ability to monitor the nucleus deformation synchronized with the electrical signal delivered to the transducer for the cyclic deformation (Fig. 1(c)), preserving the culture environment throughout the experiment.

The nucleus deformation was observed using a 50× objective lens (CFI60-2; Nikon Engineering Co., Ltd.) placed beneath the culture dish and recorded typically with a 30 ms exposure time. A trigger signal synchronized with the electrical signal for deformation was sent to the CMOS camera (C13440-20CU; Hamamatsu Photonics K.K., Japan), initiating the image acquisition. However, the start of exposure is delayed by 38.96 µs following the external trigger signal. A slight time delay is also expected because of propagation of the ultrasonic wave toward the nucleus. Therefore, the overall time delay caused throughout the experimental system was corrected by the phase delay observed in the cyclic oscillation of the silicon cantilever which is nearly perfect elastic material (Extended Data Fig. 2(c)). The phase lag of the cantilever vibration was measured ten times, resulting in -0.060±0.025 rad; this value was used as the reference phase lag for a perfect elastic material in our experimental system.

Thanks to the success of accurately evaluating the cantilever bending vibration, the phase lag in the nucleus deformation was evaluated by monitoring the mean brightness of pixels constituting the curved elements at the nucleus envelope boundary, which reflects the in-plane vibration of the nucleus. We processed obtained images as follows using Image J ver. 1.54p; (i) enhanced contrast, (ii) filter-mean with 2 pixels, and (iii) enhanced contrast again. The phase-lag analysis was conducted several times using different boundary elements for each cyclic-deformation experiment, and the average phase-lag value was determined.

The in-plane strain of nucleus was evaluated through the change in the width divided by that before deformation (Fig. 1(d)).

The typical force used for deforming the nucleus mechanical properties was 1.07 µN (or 1.19 kPa).

### RNA sequencing (RNA-seq) for single cells

We conducted the ARF-microscopy based mechanical stimulation (∼ 2 min) for several cells and prepared nucleus-stimulated and unstimulated (control) cells in the same culture dish. From each group, five nuclei were individually aspirated using a ∼10-*μ*m diameter microcapillary and transferred into separate tubes containing 10×lysis buffer (635013; Takara Bio Inc., Japan) and recombinant RNase inhibitor (2313A; Takara Bio Inc.). For the single-nucleus sampling, we used the Single Cellome™ System (SS2000; Yokogawa Electric Corporation, Japan). For the RNA-seq analysis, four tubes from nucleus-stimulated cells and two tubes from control cells were prepared, each containing five nuclei, and processed as follows. RNA libraries were first generated using the SMART-Seq mRNA protocol (Takara Bio Inc.) with the first-strand cDNA synthesis followed by PCR amplification for 18 cycles. Libraries were constructed with the Nextera XT DNA Library Prep Kit (Illumina K.K., Japan) according to the manufacturer’s instructions, and their quality was assessed with the BioAnalyzer High Sensitivity DNA Assay (Agilent Technologies, USA). Sequencing was performed on an Illumina platform to generate paired-end reads (**Source Data Fig. 2(g)_No.1**). Raw reads were processed using the DRAGEN Bio-IT Platform v4.3.6 (Illumina K.K., Japan). Reads were aligned to the human reference genome GRCh38 (GENCODE release 39), and expression levels were quantified as counts and TPM. Functional annotations were obtained from Ensembl, Gene Ontology, and Reactome databases.

Differential expression analysis was performed using the DESeq2 package (version 1.48.2) in R (version 4.5.1). Genes with zero counts in any of the six samples were excluded. Because nearly only nuclei were collected, mitochondrial genes were excluded from the analysis. Log2 fold changes were estimated using the apeglm shrinkage method implemented in DESeq2. P-values were adjusted for multiple testing using the Benjamini-Hochberg method to calculate the false discovery rate (FDR) (**Source Data Fig. 2(g)_No.2**). In the volcano plot, the horizontal axis represents the log2(fold change) between the nucleus-stimulated group and the control group, and the vertical axis represents the -log_10_(P value). Genes with an FDR value < 0.05 and |log2(fold change)| > 2 were considered significantly differentially expressed.

## Acknowledgements

We thank Mr. Shogo Kita, Mr. Yuta Terui, and Mr. Tomohiro Kaneko of Yokogawa Electric Corporation for guidance on the single-cell aspiration technique with Single Cellome™ System (SS2000). This work was supported by JSPS KAKENHI Grant No. JP24H00045.

## Extended Data Figures

**Extended Data Figure 1.**
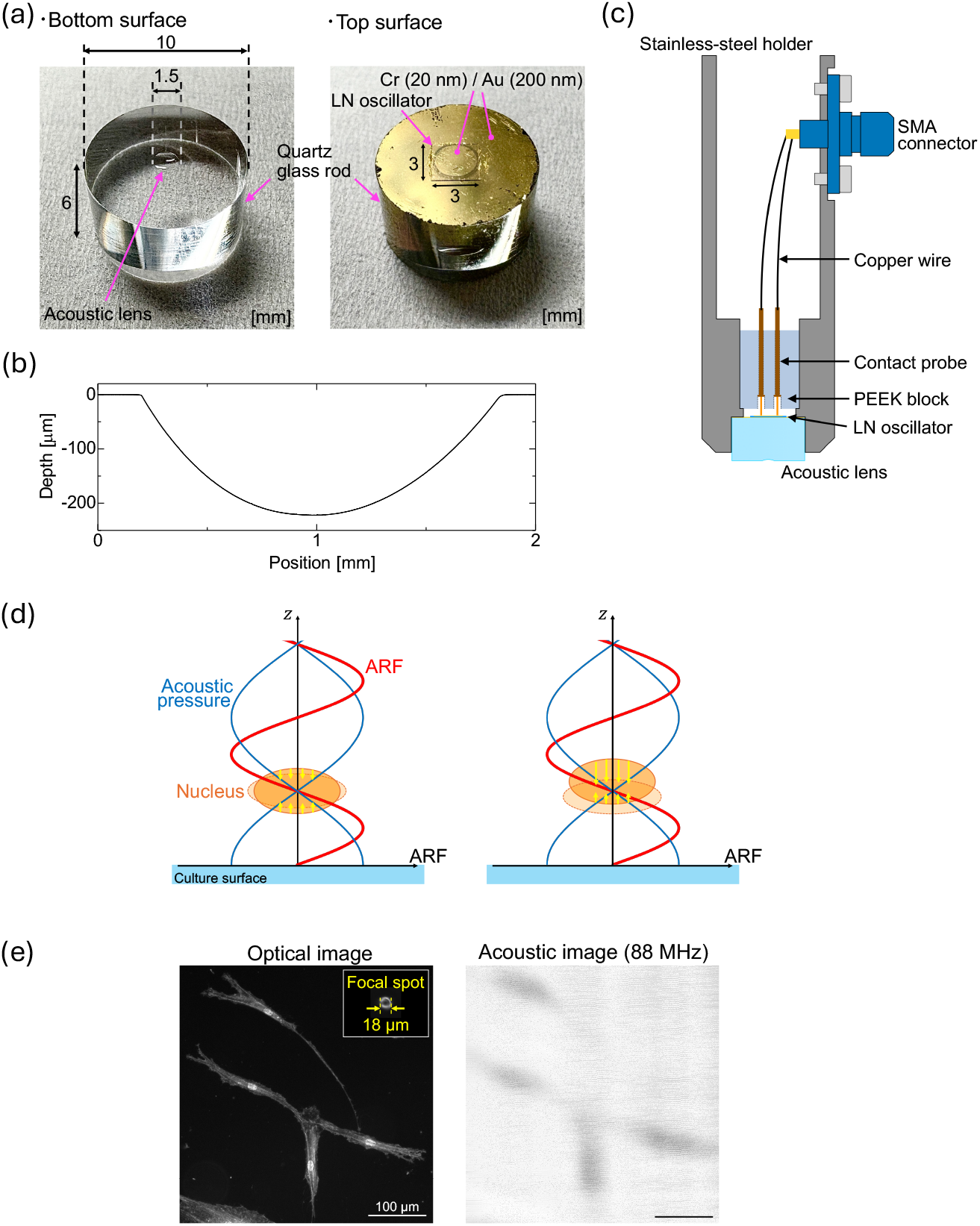
(a) The appearance of the transducer of the bottom surface with acoustic lens before depositing the electrode (left) and the top surface after the LN oscillator has been attached. The acoustic lens was fabricated by pressing a 3-mm diameter tungsten carbide sphere against the surface. (b) Cross-section surface profile of the acoustic lens measured with a stylus profilometer, confirming the smooth surface. (c) Schematic of the cross-section of the laboratory made acoustic probe. (d) Schematics of the explanation of the relationships between acoustic pressure, ARF, and nucleus movement depending on the nucleus position. Yellow arrows represent the ARF acting on the nucleus. (e) The optical image obtained by the dark-field microscopy (left) and the acoustic-absorption image at 88 MHz (right) obtained by the laboratory-made acoustic probe of human MSCs. The inset in the left figure shows the localized alterations of the plastic surface at the focal point due to ultrasonic friction.

**Extended Data Figure 2.**
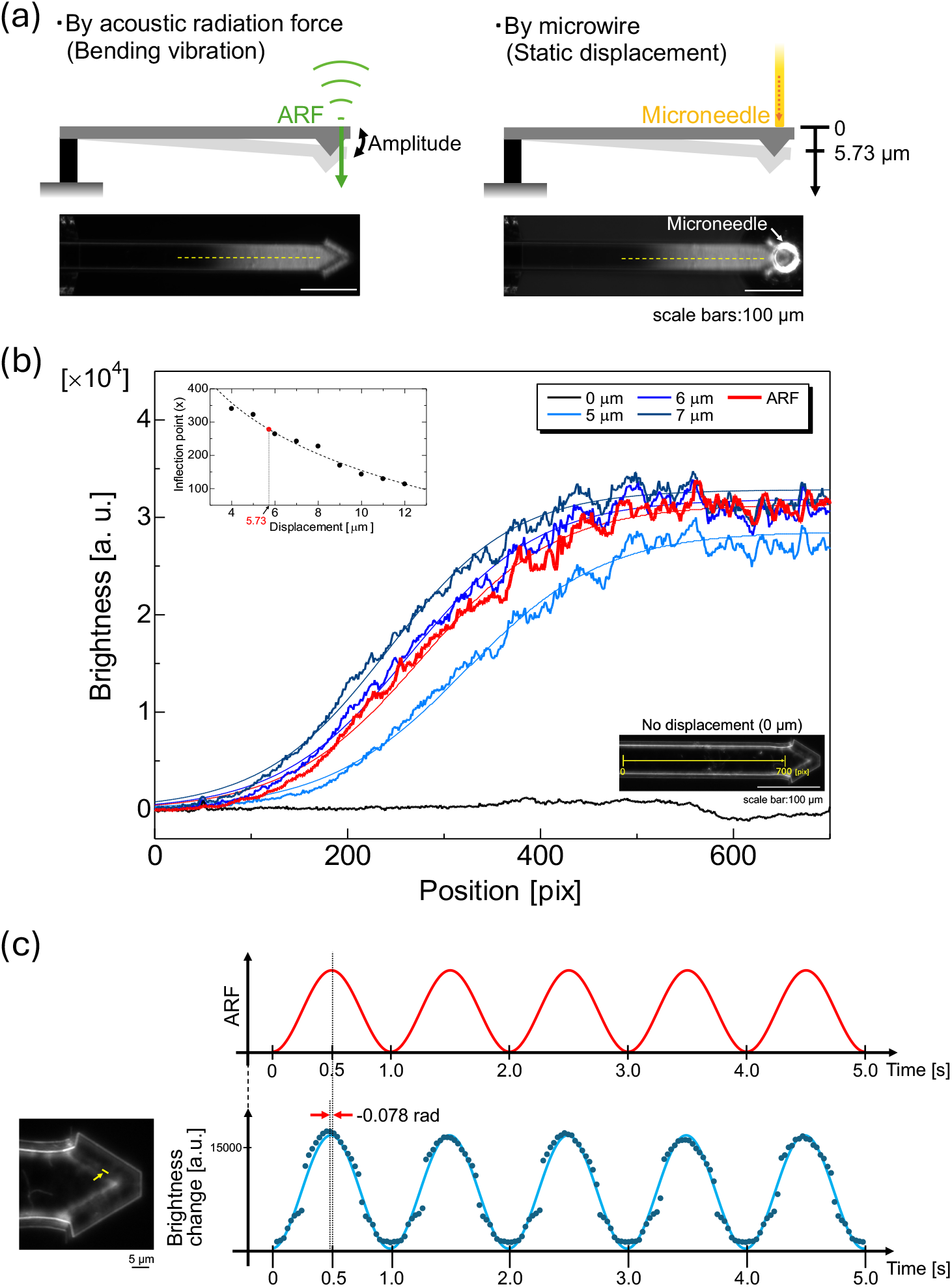
(a) Schematics of deformation of a silicon cantilever induced by ARF and microneedle indentation test. (b) The brightness changes along the cantilever (thick lines) and corresponding fitted sigmoid curves (thin lines). Each plot was transformed to set the zero-pixel value to zero. The inset represents the inflection point at each fitting sigmoid curve, from which the deformation amplitude of bending vibration was determined. (c) ARF signal (red) and changes in the average brightness of pixels (blue plots) constituting the elements near the tip of the cantilever indicated by the yellow line in the left image. The blue solid line indicates the fitted sinusoidal function.

