## Supplementary Note, Supplementary Table, Supplementary References, Descriptions of Supplementary Movies for "Focused acoustic-radiation-force microscopy for live-cell selective nucleus deformation and nucleus viscoelasticity measurement"

Supplementary Note  Page S2-S4
Supplementary Table 1  Page S5
Supplementary Reference  Page S6-S7
Descriptions of Supplementary Movies S8

**Supplementary Note: Estimation of temperature increase in nucleus**

It is well known that energy flow of an acoustic plane wave (absolute value of the acoustic Poynting vector) is estimated by $I = P_{ac}^2/(2\rho c)$, which represents the energy inflow per unit area per unit time. Thus, a simple estimation for the acoustic power $W_{ac}$ injected into a nucleus with radius $a$ is made as follows

$$W_{ac} = \frac{\pi a^2 P_{ac}^2}{2\rho c}. \tag{1}$$

Here, $P_{ac}$, $\rho$ and $c$ represent the acoustic pressure amplitude, mass density and sound velocity of the culture medium, respectively. Because a part of this energy is absorbed in the nucleus, one may expect a significant temperature increase over time. However, nucleus is surrounded by cytoplasm, which dissipates the heat from the nucleus.

For investigating this effect, we analyzed the temperature rise in nucleus subjected to the ultrasonic heating. First, we evaluated the ultrasonic attenuation coefficient $\alpha_n$ inside nucleus by comparing the amplitudes of the ultrasound wave reflected at the dish surface in the presence and absence of nucleus along the propagation path. (The amplitude was actually decreased as shown in the inlet of Fig. 1(b)). Letting $\tilde{A}$ and $d$ denote their amplitude ratio and the nucleus thickness, respectively, $\alpha_n$ is given by $\alpha_n = -\ln\tilde{A}/2d$, noting that the ultrasonic wave passes through the nucleus twice. Because the attenuation coefficient for the energy flow equals $2\alpha_n$, the adsorbed energy per unit time within the nucleus is estimated by $W_{nuc} = 2W_{ac}(1 - \exp(-2\alpha_n d))/2$. The former factor 2 appears because of heating by incident and reflected waves and the latter factor 2 by the cyclic heating due to modulation.

We consider that this ultrasonic heating occurs uniformly within the nucleus and solved the heat conduction equation under a spherical symmetry

$$\left\{\frac{1}{r^2}\frac{\partial}{\partial r}\left(r^2\frac{\partial T}{\partial r}\right)\right\} + \frac{W}{V\kappa} = 0\,, \tag{2}$$

where

$$W = \begin{cases} W_{nuc} & (r \le a) \\ 0 & (r > a) \end{cases}. \tag{3}$$

Here, $r$, $T$, $V$ and $\kappa$ denote radius, temperature, volume and thermal conductivity, respectively.

Since $W = 0$ when $r > a$,

$$\frac{\partial}{\partial r}\left(r^2\frac{\partial T}{\partial r}\right) = 0, \tag{4}$$

we obtain

$$r^2 \frac{\partial T}{\partial r} = C_1 \quad (C_1 : constant). \tag{5}$$

Therefore,

$$T = C_2 - \frac{C_1}{r}. \tag{6}$$

By redefining the constant, for $r > a$, that is, the cytoplasm temperature, $T_c$, is given as follows

$$T_c = C_1 + \frac{C_2}{r}. \tag{7}$$

Setting the temperature at $r \to \infty$ as $T_c = T_0$,

$$T_c = T_0 + \frac{C_2}{r}. \tag{8}$$

Then, the heat flow rate through the surface of the sphere is given by

$$4\pi a^2 \left(-\kappa_c \frac{\partial T_c}{\partial r}\right)_{|r=a} = 4\pi \kappa_c C_2, \tag{9}$$

where $\kappa_c$ describes thermal conductivity of cytoplasm. As this equals the heat generation within the nucleus, $W_{nuc}$, it follows that

$$C_2 = \frac{W_{nuc}}{4\pi \kappa_c}. \tag{10}$$

We thus obtain $T_c$ as

$$T_c = T_0 + \frac{W_{nuc}}{4\pi \kappa_c} \frac{1}{r}. \tag{11}$$

While $W = W_{nuc}$ for $r < a$, we obtain

$$\frac{\partial}{\partial r}\left(r^2 \frac{\partial T}{\partial r}\right) = -\frac{W_{nuc}}{\kappa_{nuc} V_{nuc}} r^2. \tag{13}$$

Therefore,

$$\frac{\partial T}{\partial r} = -\frac{1}{3} \frac{W_{nuc}}{\kappa_{nuc} V_{nuc}} r + \frac{C_3}{r^2}. \quad (C_3 : constant) \tag{14}$$

Because $\partial T / \partial r$ remains finite at $r = 0$, we have $C_3 = 0$. Thus, the temperature in the nucleus, $T_{nuc}$, is given as follows

$$T_{nuc} = -\frac{1}{6} \frac{W_{nuc}}{\kappa_{nuc} V_{nuc}} r^2 + C_4. \quad (C_4 : constant) \tag{15}$$

Since $T_{nuc} = T_c$ at $r = a$, $C_4$ is given by

$$C_4 = T_0 + \frac{W_{nuc}}{4\pi\kappa_w}\frac{1}{a} + \frac{1}{6}\frac{W_{nuc}}{\kappa_{nuc}V_{nuc}}a^2. \qquad (16)$$

Hence,

$$T_{nuc} = T_0 + \frac{W_{nuc}}{4\pi\kappa_w}\frac{1}{a} + \frac{1}{6}\frac{W_{nuc}}{\kappa_{nuc}V_{nuc}}(a^2 - r^2). \qquad (17)$$

Therefore, the temperature at the center of the nucleus is described as

$$T_{nuc} = T_0 + \frac{W_{nuc}}{8\pi a}\left(\frac{2}{\kappa_c} + \frac{1}{\kappa_{nuc}}\right). \qquad (18)$$

Typical values for parameters in our experiments ($P_{ac}$= 4.0 MPa, $a$=5 mm, $\tilde{A}$=0.92, and $d$=5 mm, $\kappa_c$=$\kappa_{nuc}$=0. 6 $Wm^{-1}K^{-1}$) yield a temperature increase approximately 1 ºC even at the center of the nucleus (the hottest spot).

In fact, to assess the thermal effect, we placed a needle thermocouple in the culture medium and continuously applied focused ultrasound to its tip for ~10 min. However, no measurable temperature change was detected within the sensitivity limit (0.1 ºC) of the thermometer.

### Supplementary Table 1: A summary of methods for measuring nucleus mechanical properties and for its mechanical stimulation

| methods | non-invasiveness | feasibility of measurements confined to a single cell nucleus without nucleus isolation | feasibility of a long-time measurement for nucleus of the same living cell | notes |
|---|---|---|---|---|
| atomic-force-microscopy [1-5] | × | ○ | × | Physical contact between the cantilever and the cell can alter its structure, including the cytoskeleton and the cell membrane, or cause damage. |
| micropipette [5-7] | × | ○ | × | Physical contact between the pipette tip and the cell can alter its structure, including the cytoskeleton and the cell membrane, or cause damage. |
| micromanipulator [8,9] | × | ○ | × | Physical contact between the probe and the cell can alter its structure, including the cytoskeleton and the cell membrane, or cause damage. |
| magnetic tweezer [10-14] | × | ○ | × | Chemical linking between nucleus and microbeads is needed for force application. Significant thermal effects due to high-intensity magnetic fields are expected. |
| optical tweezer [15-19] | × | ○ | × | Chemical linking between nucleus and microbeads is needed for force application. Significant thermal effects due to high-intensity laser beams are expected. |
| substrate stretching method [20-23] | ○ | × | ○ | This method deforms the entire cell, including cytoplasm, making it difficult to isolate the specific mechanical contributions sensed by the nucleus. |
| acoustic tweezer [24-30] | ○ | × | ○ | High mobility of suspended cells precludes the ability to target nuclear deformation specifically. |
| ARF microscopy (present study) | ○ | ○ | ○ | The only technique that can non-invasively and selectively apply mechanical stimulation to a single nucleus in a living cell while enabling long-term measurement of its viscoelastic properties. |

**Descriptions of Supplementary Movies**

**Supplementary Movie S1:** Nuclei deformations at 1 Hz monitored using the dark-field microscope. Note that although the four movies are combined into one, they were measured individually.

**Supplementary Movie S2**: A 1-Hz deformation can be induced only in the target nucleus, indicated by a red arrow, within a wide field of view.
